# Physiological stress response of koala joeys to visitors

**DOI:** 10.1101/2023.10.09.561560

**Authors:** Harsh K. Pahuja, Izzy Bee, Ali Bee, Edward J. Narayan

## Abstract

Koala (*Phascolarctos cinereus*) joey rescues are increasing over the years, and rehabilitation of a joey requires extensive care, close proximity and handling by humans. These novel environments are likely to present a suite of biotic and abiotic stressors during rehabilitation. In this study, we longitudinally monitored the faecal cortisol metabolites (FCMs) of three koala joeys within the context of potential stressors at the Magnetic Island Koala Hospital, Queensland, Australia. A total of 92 faecal samples were analysed for FCMs using a polyclonal R4866 cortisol enzyme-immunoassay which has been previously validated in koalas. The iterative baseline method was used to establish FCM profiles of all individuals, and to identify significant peaks in FCM concentrations. Visitor events were identified and confirmed as an acute stressor based on the FCM profiles of the koala joeys. All three koala joeys elicited a significant rise in FCM concentrations after each visitor encounter. To our knowledge, this is the first study to report on the acute stress response of koala joeys to visitors. We recommend that visitor encounters be kept to a minimum, and perhaps avoided altogether especially for joeys that are being rehabilitated to be released back into the wild.

## Introduction

The Koala (*Phascolarctos cinereus*) is an iconic marsupial endemic to Australia (Woinarski & Burbidge, 2020). Retrospective studies assessing the trends of koala admissions to wildlife hospitals highlight that the clinical outcome for the majority of these koalas is that they either naturally succumb to their injuries and/or diseases, or they need to be humanely euthanised due to their critical condition (Burton & Tribe, 2016; Gonzalez-Astudillo et al., 2017; Griffith et al., 2013; Taylor-Brown et al., 2019). Increased mortality of adult female koalas is likely to cause more rescues of young orphans because during the early stages of their life, joeys are sustained by their mothers. In the study by Taylor-Brown et al. (2019), koala joeys formed the second largest group (after possums) that were rescued as orphans. Furthermore, as highlighted by Pahuja and Narayan (2023a), koala joey rescues are also increasing slightly over the years. Koala joeys are born in an altricial state, which is followed by a ∼6-month period of pouch occupancy (Gipp, 2004). During this extensive dependant period, maternal milk, eucalyptus and ‘pap’ constitute the diet of joeys (Barker et al. 2013; Morris et al. 2016). When orphaned, of course, this process needs to be replicated by humans and requires extensive care, close proximity and handling. Being raised by a human in captivity is likely to serve as a ‘stressor’ for the koala joey because of the novel environment.

A stressor is commonly defined as any noxious stimuli that could potentially cause an imbalance in the animal’s homeostatic system (Moberg & Mench, 2000). When encountered with a stressor, the body elicits an array of responses; one of which is the neuroendocrine activation of the hypothalamo-pituitary-adrenal (HPA) axis (Sheriff et al., 2011). This results in the secretion of glucocorticoids (GCs) which have a range of effects (e.g. energy mobilisation for fight-or-flight response) that allow the animal to achieve homeostasis i.e. cope with the stressor (Moberg & Mench, 2000). As a result, GCs are often referred to as the stress hormone (Cope et al., 2022), and short-term elevation of GCs are indicative of acute stress (Hogan et al., 2012). Prolonged GC elevation can have deleterious impact on an animal’s health given the diverse biological effects of GCs on various body systems, and is thus indicative of chronic stress (Hogan et al., 2012; Narayan et al., 2012). GCs (or its metabolites) can be measured in a range of sample types, such as blood, faeces and fur. However, given the ease of collection and minimal handling associated with collecting scat samples, monitoring faecal glucocorticoid metabolites (FGM)s is the most commonly employed method in studies on Australian marsupials (Hing et al., 2014). Cortisol is the principal GC produced by koalas (Narayan et al., 2013) and is increasingly acknowledged as a practical endocrine indicator of stress (Hing et al., 2014; Narayan & Williams, 2016). Whilst there have been a number of studies investigating the physiological stress response of marsupials (Cope et al., 2022; Hing et al., 2017; Hogan et al., 2011; Hogan et al., 2012; Narayan et al., 2012), including koalas (Charalambous et al., 2021; Pahuja & Narayan, 2023b; Webster et al., 2017), there is a paucity of literature on the capacity of koalas to respond to the stressors of captivity. In order to improve the rehabilitation outcomes and ensure the welfare of koalas in captivity, it is imperative to identify the potential stressors of captivity and understand how these animals respond to it.

In the present study, we monitored the faecal cortisol metabolites (FCM)s of three orphaned koala joeys undergoing rehabilitation over a period of one-month before release. During this period, we also monitored potential stressors faced by the joeys during rehabilitation. Of particular interest then was to examine whether koala joeys elicit a physiological stress response to these stressors. We hypothesised that stressors are likely to cause a physiological change in the joeys which will be reflected and capture as elevated FCMs. The primary aim of this study was to expand the current knowledge on koala stress biology by monitoring FCMs within the context of potential stressors. To our knowledge, this is the first study to report on the physiological stress response of koala joeys to potential stressors.

## Material and Methods

### Animal Ethics

All procedures in this research were performed in accordance with the relevant guidelines and regulations set by The University of Queensland Animal Ethics Committee: Native or Exotic Wildlife and Marine Animals (NEWMA) (approval number 2022/AE000035).

### Study Site and Animals

This research study was conducted in collaboration with the Magnetic Island Koala Hospital. All koala joeys were housed at 3 Corica Crescent, Horseshoe Bay, QLD, Australia 4819 (GPS coordinates: 19°7′26.06″S, 146°51′34.06″E). A total of 3 orphaned koala joeys [1 male (M1) and 2 females (F1 and F2)] were studied based on whichever was currently in care during the period of data collection (2^nd^ April 2022 to 2^nd^ May 2022). All koala joeys were housed individually in a 6 x 6 x 3.5 (L x W x H) meter enclosure donated by the Glencore Australia, Copper Refineries Pty. Ltd. The enclosure was designed with a cement slab floor, insulated roof, snake proof mesh and trim deck metal, and furnished with logs, branches, ropes and polyvinyl pipes of varying dimensions. The joeys were provided with *ad libitum* water and feed. The feed consisted of a mix of various species of *Eucalyptus* leaves freshly cut and sourced from a local *Eucalyptus* plantation. The *Eucalyptus* leaves were changed daily and mainly consisted of (but not limited to) river red gum (*E. camaldulensis*), swamp mahogany (*E. robusta*), red mahogany (*E. resinifera*), mountain mahogany (E. notabilis) and tallow (*E. microcorys*) The age for all koala joeys during data collection was estimated by the veterinarian to be between 7-9 months based on tooth wear and body weight. No joeys had any history of disease, and remained clinically healthy throughout the study period as reported by the veterinarian.

### Data collection (Faecal sample collection and recording stressors)

In order to evaluate daily changes in FCM concentrations, fresh faeces were collected routinely from all three koala joeys daily, over a one-month study period. One faecal sample (comprising on average of 3-5 faecal pellets) per day per animal was collected during routine cleaning of the enclosure (0900h to 1100 h), with minimal disturbance to the animal. Samples were collected by hand, using a new pair of gloves, and immediately transferred into a resealable polyethylene ZipLock© bag and labelled with the animal’s name, date and the time of sample collection. All faeces were removed from each enclosure at the end of each day so that we could ensure fresh faecal sample collection on the following day. The samples were then stored at −20°C until they were sent to the laboratory on ice via overnight freight. The faecal samples were transited on ice via overnight freight and upon delivery, they were immediately refrozen at −20°C until processing.

The staff at the Magnetic Island Koala Hospital also observed and recorded any potential stressors encountered by the joeys. A range of stressors have been identified for marsupials in captivity [see Hing et al. (2014), Hogan et al. (2012), Charalambous et al. (2021)], including (but not limited to) maintenance activities in and around the enclosure, moving animals to a different enclosure, visitor encounters, loud noises, extreme environmental events, handling for routine health checks and so on. This non-exhaustive list of potential stressors was used as a reference to observe and record potential stressors that could be affecting the koala joeys.

### Faecal sample processing and extraction

Faecal sample processing and FCM extraction followed the procedures previously described for koalas (both, adults and joeys) (Charalambous et al., 2021; Narayan et al., 2013; Pahuja & Narayan, 2023b) Briefly, frozen faecal samples were weighed and then dried in an oven at 65°C for a 24 h period, or until completely dried. Once completely dry, the samples were ground into a fine powder using a mortar and pestle. Analytical grade ethanol (10% v/v) was used to clean the mortar and pestle between each individual sample processing to avoid cross-contamination. The ground-up powder was sifted through a fine mesh strainer to remove all coarse particles. A 0.2 ± 0.001 g sample of sifted product was weighed out into a labelled Eppendorf tube and then stored in a −20°C freezer until use.

For hormone extraction, samples were removed from the −20°C freezer and 2 mL of 90% (v/v) analytical grade ethanol solution was added to the Eppendorf tube. Tubes were vortexed at medium-high speed (5 min, 5000g) on an Eppendorf mini-spin centrifuge for a minimum of 30 s to thoroughly mix the solution. Tubes were then placed into a water bath (>60°C) for 20 min to allow hormones to dissolve in the solution. While in the bath, tubes were gently manually shaken to ensure faeces remained submerged in ethanol and did not spill over the top of the tube.

After 30 min, the supernatant was transferred into an Eppendorf tube, closed and then centrifuged at 3000g for 5 min to separate any solid faecal residue. If the sample looked turbid or had suspended particles, tubes were centrifuged again for 5 min and then pipetted into a new and clean labelled Eppendorf tube. Following this, for the polyclonal cortisol antibody assay (R4866; Late C. J. Munro, University of California, Davis), 0.6 mL of the aqueous solution was aliquoted into a new and clean labelled Eppendorf tube. The tubes were kept in a laminar flow chamber for 24 h to allow enough time for the ethanol to completely evaporate. Once ethanol was completely evaporated, 1 mL of assay buffer (39 mM NaH_2_PO_4_·H_2_O, 61 mM NaHPO_4_, 15 mM NaCl and 0.1% bovine serum albumin, pH 7.0) was added to each tube. The tubes were vortexed at medium-high speed (5 min, 3000g) in an Eppendorf mini-spin centrifuge for 30 s, and then centrifuged at 10000g for 10 min. Following this, 850 µL of supernatant was pipetted into a new and clean labelled Eppendorf tube, ensuring any solid remnants in the solution were avoided.

### Enzyme-immunoassay (EIA)

All consumables for the EIA were obtained from Sigma-Aldrich, The R4866 EIA has previously been validated in koalas using physiological (ACTH challenge), biological and laboratory validation techniques, and is routinely used for monitoring physiological stress in koalas (Charalambous et al., 2021; Narayan, 2019; Narayan & Vanderneut, 2019; Narayan et al., 2013; Webster et al., 2017). In addition, this assay has also been validated in the laboratory for monitoring FCMs in koala joeys (Pahuja & Narayan, 2023b). Glucocorticoid concentrations for faecal extract using the R4866 assay were determined using a polyclonal anti-cortisol antiserum diluted to 1:15 000, horseradish peroxidase (HRP) conjugated cortisol 1:80 000 and cortisol standards (1.56–400 pg/well). Cross reactivity of the R4866 anti-cortisol antiserum is reported as 100% with cortisol and less than 10% with other steroids (with prednisolone 9.9%, prednisone 6.3%, cortisone 5% and < 1% with corticosterone) tested (Munro & Stabenfeldt, 1985). Sample extracts were assayed in duplicate on Nunc Maxisorp™ plates (96 wells) (Sigma Aldrich, Sydney, Australia). Plates were coated with 50 µL cortisol antibody diluted to the appropriate concentration in a coating buffer (50 mmol/L bicarbonate buffer, pH 9.6), and left to stand and incubate for a minimum 12 h in a fridge at 4°C. The plates were washed the next day using an automated plate washer (ELx50, BioTek™, Sursee, Switzerland) with phosphate-buffered saline containing 0.5 mL/L Tween 20 to rinse away any unbound antibody.

Stocks of standards, faecal extracts and horseradish peroxidase labels were diluted to the appropriate concentration in assay buffer. The extraction dilution rates (1:2) were based on the concentrations of pooled samples that resulted in 50% binding in a previous study monitoring FCMs in orphaned koala joeys (Pahuja & Narayan, 2023b). For each assay, 50 µL of cortisol standard and diluted faecal extract was added to each well based on the plate map.

Immediately following this, 50 µL of HRP label was added to the wells. Plates were covered and incubated at room temperature for 2 h. Following a wash, 50 µL of substrate buffer (0.01% tetramethylbenzidine and 0.004% H_2_O_2_ in 0.1 M acetate citrate acid buffer, pH 6.0) was then added to generate a colour change. Colour reaction was halted after 15 min using 50 µL of stop solution (50 μL of 0.5 mol/L H_2_SO_4_), and the plates were read primarily at 450 nm and secondarily at 630 nm on an ELx800 (BioTek) microplate reader. The sensitivity of the faecal cortisol FCM was calculated as the value 2 standard deviations from the mean response of the blank samples. The sensitivity of the faecal cortisol metabolite EIA was 1.2 ± 0.2 pg well^-1^ (n = 3). Intra- and inter-assay coefficients of variation (CV) were determined by calculating the mean of CV within a plate (for intra-assay) and between the plates (for inter-assay) assayed during analysis. The intra- and inter-assay coefficients of variation were 4.26 ± 1.10 % (n = 3) and 6.42 ± 3.71 % (n = 3) respectively.

### Data processing and analysis

All FCM concentrations were expressed as (ng/g) on the basis of net dry faeces weight. Analysis of monthly profiles of FCM secretion involved establishing a baseline level in order for significant peaks to be identified. FCM concentrations from individual koala joeys were analysed using an iterative baseline method (Brown et al., 1994). In this method, the baseline is calculated through an iterative process, excluding concentrations greater than ‘mean + 2 SD’. In successive iterations, all concentrations greater than ‘mean + 2 SD’ were excluded until no points fall above this threshold. The mean iterative baseline was the mean of the remaining concentrations at the completion of the iterative process. Concentrations that fall above the threshold (mean + 2 SD) of the final iteration were classified as significant peaks. The ‘mean + 2 SD’ of the final iteration represents the ‘cut-off’ (i.e. baseline threshold) value and all concentrations above this value were considered to be an indicative of a significant acute stress response. Several measures were calculated to summarize faecal hormone profiles for each individual koala joey : (1) overall mean of all samples for the collection period, (2) baseline mean that excluded all values greater than ‘mean + 2 SD’ of the final iteration, and (3) peak mean that only included all values > 2 SD above the baseline mean. These values are highlighted in Table 1. The iterative baseline calculation can be performed using the R package hormLong derived by Fanson and Fanson (2015) for large datasets and/or when monitoring multiple hormonal profiles. However, given the nature and scope of the current study, these calculations were performed in Microsoft Excel 2021©.

**Table 1.**
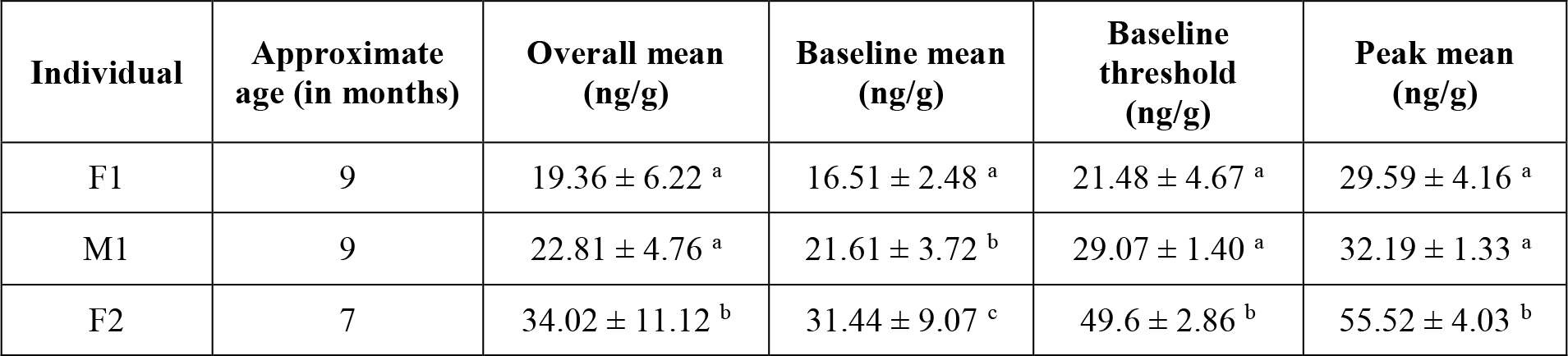
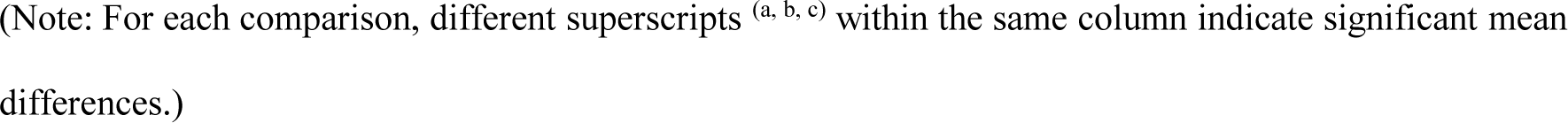
Iterative baseline method representing the overall mean, baseline mean, baseline threshold and peak mean for each individual koala joey involved in the study.

| Individual | Approximate age (in months) | Overall mean (ng/g) | Baseline mean (ng/g) | Baseline threshold (ng/g) | Peak mean (ng/g) |
| --- | --- | --- | --- | --- | --- |
| F1 | 9 | $19.36 \pm 6.22^a$ | $16.51 \pm 2.48^a$ | $21.48 \pm 4.67^a$ | $29.59 \pm 4.16^a$ |
| M1 | 9 | $22.81 \pm 4.76^a$ | $21.61 \pm 3.72^b$ | $29.07 \pm 1.40^a$ | $32.19 \pm 1.33^a$ |
| F2 | 7 | $34.02 \pm 11.12^b$ | $31.44 \pm 9.07^c$ | $49.6 \pm 2.86^b$ | $55.52 \pm 4.03^b$ |

(Note: For each comparison, different superscripts <sup>(a, b, c)</sup> within the same column indicate significant mean differences.)

### Statistical analysis

Statistical analysis was performed in GraphPad Prism version 9.3.1. The parametric assumptions of normality and homoscedasticity were tested and a single factor ANOVA, followed by Tukey’s multiple comparison test, was run to determine whether overall mean, baseline mean, baseline threshold and peak mean FCM concentrations differed (*P* < 0.05) among individuals.

## Results

### Identification of stressors

The staff at the Magnetic Island Koala Hospital identified visitors as the only potential stressor for the three koala joeys undergoing rehabilitation. Visitors mainly included potential donors and their associated members and/or media, in an attempt to raise funding for improving the conservation of koalas. A visitor event could be described as photographing the joeys from a safe distance from inside the enclosure and/or media interviews to raise awareness. Visitors were always accompanied by hospital staff and followed a strict no-contact policy. Visitors were confirmed as a stressor based on the FCM profiles of all three koala joeys. These results are presented in detail below.

### FCM profiles and acute stress response of koala joeys to visitors

The daily FCM concentrations of the three koala joeys (Fig. 1) ranged from 10.41 to 33.27 ng/g dry faeces for individual F1, 14.42 to 33.65 ng/g dry faeces for individual M1 and 15.41 to 59.82 ng/g dry faeces for individual F2. There were significant differences in the overall mean (F_2,88_ = 7.689, p < 0.05), baseline mean (F_2,75_ = 13.21, p < 0.05), baseline threshold (F_2,9_ = 1.104, p < 0.05) and peak mean (F_2,9_ = 0.503, p < 0.05) FCM concentrations between the three koala joeys over the one-month sampling period (Table 1). The overall mean FCM concentrations of F1 (19.36 ± 6.22 ng/g dry faeces) and M1 (22.81 ± 4.76 ng/g dry faeces) were significantly lower (p < 0.05 for both comparisons) relative to F2 (34.02 ± 11.12 ng/g dry faeces), but did not differ significantly amongst each other (p = 0.24) (Table 1). The iterative baseline mean FCM concentrations of all three koala joeys were significantly different from each other (p < 0.05 for all comparisons) (Table 1). The baseline threshold FCM concentrations of F1 (21.48 ± 4.67 ng/g dry faeces) and M1 (29.07 ± 1.40 ng/g dry faeces) were significantly lower (p < 0.05 for both comparisons) relative to F2 (49.6 ± 2.86 ng/g dry faeces), but did not differ significantly amongst each other (p = 0.09) (Table 1). The peak mean FCM concentrations of F1 (29.59 ± 4.16 ng/g dry faeces) and M1 (32.19 ± 1.33 ng/g dry faeces) were significantly lower (p < 0.05 for both comparisons) relative to F2 (55.52 ± 4.03 ng/g dry faeces), but did not differ significantly amongst each other (p = 0.78) (Table 1). There were no consistent patterns in FCM profiles of koala joeys in relation to their age and/or sex (Fig. 1).

**Fig. 1.**
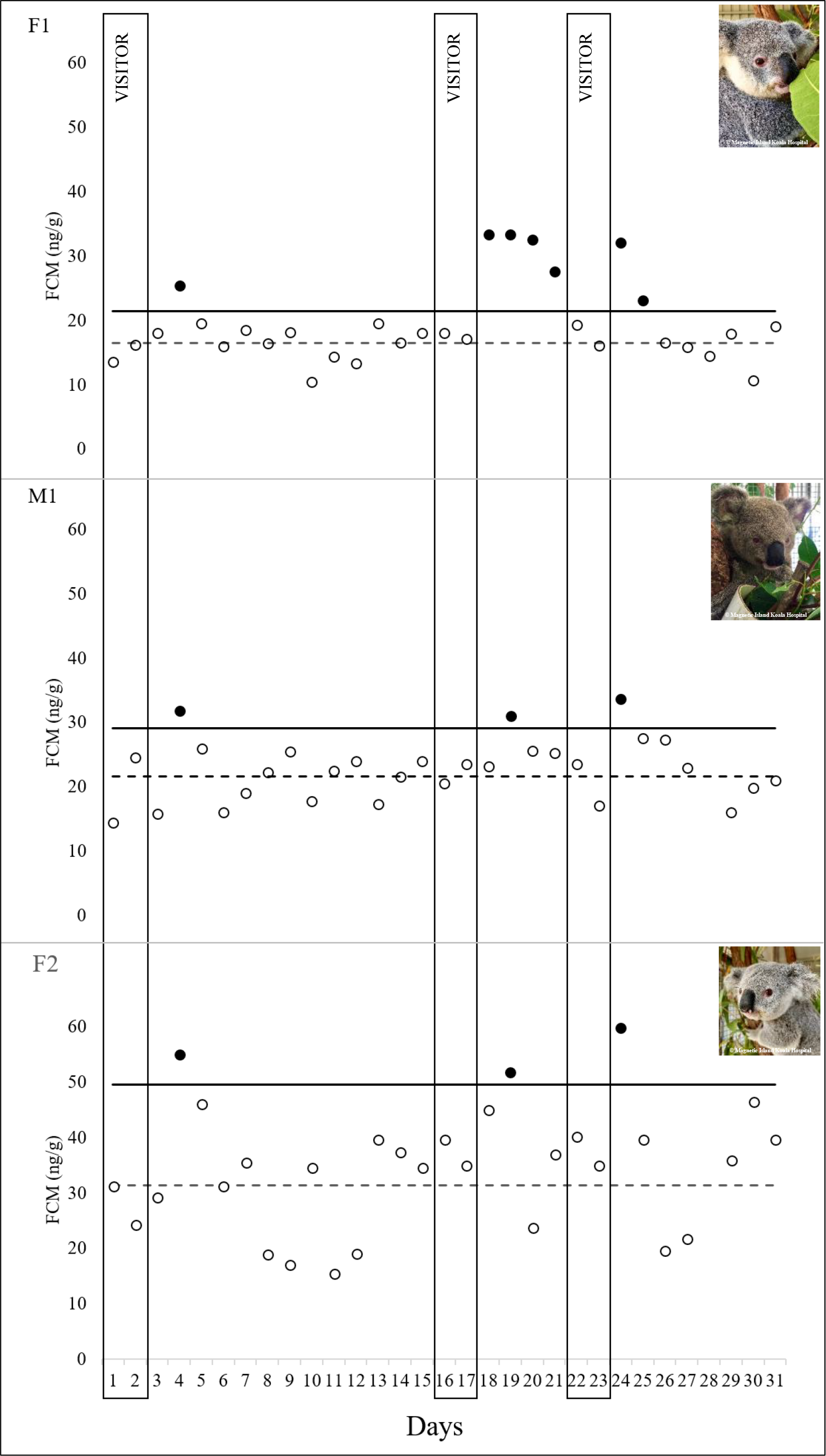
FCM profiles of koala joeys undergoing rehabilitation at the Magnetic Island Koala Hospital. All open circles illustrate baseline FCM concentrations, whereas closed circles illustrate significant peaks in FCM concentrations as identified by the iterative baseline method. Vertical boxes delineate visitor events as identified by the staff at Magnetic Island Koala Hospital. Solid horizontal line represents the baseline threshold value. Dashed horizontal line represents the mean iterative baseline.

All koala joeys displayed a significant increase (above baseline threshold) in FCM concentrations within 1-3 days after each of visitor event (Fig. 1). After the 1^st^ visitor event (Days 1-2), all koala joeys exhibited a significant peak in FCM concentrations (Day 4) followed by an immediate return to baseline concentrations. The FCM concentrations of all koala joeys remained within the baseline threshold limit (Days 5-14) until the next visitor event (Days 15-16). Following the 2^nd^ visitor event, all koala joeys exhibited a significant peak in FCM concentrations (F1: Day 18; M1: Day 19; F2: Day 19) (Fig. 1). The FCM concentrations of M1 and F2 immediately returned within the baseline threshold concentrations until after the next visitor event. It is important to note that for F1, the FCM concentrations decreased consistently over the following days, but remained above the baseline threshold. Following the 3^rd^ visitor event (Days 22-23), all koala joeys exhibited a significant peak in FCM concentrations (Day 24). The FCM concentrations of M1 and F2 immediately returned within baseline threshold, however, the return to baseline was slightly delayed for F1 (Fig. 1). The FCM concentrations of all joeys remained within the baseline threshold until they were released on Day 31.

## Discussion

The primary aim of this study was to monitor FCMs in koala joeys within the context of potential stressors. As hypothesised, potentially stressful stimuli (i.e. visitors) resulted in an increase in FCM concentrations on all three encounters, across all koala joeys. These findings confirm that visitors elicit an acute stress response in koala joeys. To our knowledge, this is the first study to report on the acute stress response of koala joeys to visitors.

Individual differences in the FCM profiles of koala joeys was not an unexpected finding. These results are consistent with previous research on rescued koala joeys highlighting inter-individual differences (Pahuja & Narayan, 2023b). In the current study, the natural variability in FCM profiles is reflected in the overall mean, baseline mean, peak mean, baseline threshold and daily variation in FCM concentrations of the koala joeys. GCs are known to reflect inter- and intraindividual variation which can be mediated by several factors such as age, sex, body condition, genetics, season, habituation, temperament, personality and so on (Cockrem, 2013; Guindre-Parker, 2020). When considering baseline FCM concentrations, F2 had significantly higher baseline FCM concentrations relative to F1 and M1. As highlighted by Guindre-Parker (2020), baseline GCs are likely to be strongly correlated with energetic balance of the animal. Since F2 is the youngest animal, it is possible that the elevated baseline FCM concentrations of F2 are a result of its high energetic demands. However, it is also possible that since F2 had spent the least amount of time under rehabilitation, it was habituated to a lesser degree to the captive environment resulting in elevated baseline FCM concentrations [see (Hogan et al., 2012)]. Furthermore, as noted by the veterinarian, F2 was rejected by the mum while in care, which could also contribute to the high baseline FCM concentrations [see (Romero et al., 2009)]. Therefore, it is impossible to isolate a single factor that causes variability among FCM profiles of individuals, because it is most likely the result of synergistic interaction between numerous factors.

All three visitor events evoked a significant acute stress response in all joeys. It has been documented previously that visitors can evoke a stress response from adult koalas subjected to photography regimes (Webster et al., 2017). Although, in our study all three visitor events evoked a significant acute stress response in all joeys, the magnitude of stress response and the time taken to return to baseline concentrations differed for each individual and for each visitor event. Incorporating these findings into the reactive scope model (RSM) [as proposed by (Romero et al., 2009)] highlights new avenues about the physiological coping mechanisms of koalas. The RSM proposes that the physiological mediators associated with the stress response can be divided into four distinct ranges – (i) predictive homeostasis (ii) reactive homeostasis (iii) homeostatic failure and (iv) homeostatic overload. The predictive range encompasses physiological mediator levels that vary in response to predictable environmental change i.e. tidal, seasonal and circadian rhythm. The reactive range encompasses physiological mediator levels that vary in response to unpredictable changes i.e. response to stressors. The predictive and reactive range together form the normal reactive scope of an animal, and is essential for the day-to-day functioning of the animal. The homeostatic overload range encompasses physiological mediator levels above the normal reactive scope i.e. chronic stress. The homeostatic failure range encompasses physiological mediator levels below the normal reactive scope i.e. insufficiency to respond to stressors. The homeostatic overload and failure serve as two extremes of the RSM and are associated with detrimental pathological effects (Romero et al., 2009).

In our study, the baseline concentrations (represented by open circles in Fig. 1) can be presumed to be an indicative of the predictive homeostasis range, representing the natural rhythm in FCMs. After the 1^st^ visitor event, the FCM concentrations of all individuals increased significantly, indicating a classic activation of the HPA axis, followed by an immediate return to baseline concentrations indicating the deactivation of the HPA axis (via the negative feedback loop) once the stressor had passed (Sheriff et al., 2011). The significant increases in FCM concentrations (represented by closed circles in Fig. 1) can be presumed to be an indicative of the reactive homeostasis range wherein the transient rise in FCM concentration enables the animal to cope with the stressor and re-establish homeostasis (Wingfield, 2013). Following the 2^nd^ visitor event, the FCM concentrations of F1 were elevated for a longer duration, relative to M2 and F2 which followed a similar pattern of immediately returning to baseline concentrations. Two scenarios are possible in this case for F1 – (i) presence of a prolonged stressor i.e. visitors were present on successive days (ii) delayed termination of stress response. Both of these scenarios make the animal susceptible to enter the homeostatic overload range that can have deleterious impact on the animal (Romero et al., 2009). Since the staff at the Magnetic Island Koala Hospital confirmed the absence of visitors on successive days, the presence of prolonged stressor can be ruled out. Therefore, it could be presumed that the termination of the stress response was delayed due to weaker efficacy of the negative feedback loop. Delayed termination of stress response and its impact on the animal has been discussed thoroughly by Romero (2012) by highlighting the example of Galápagos marine iguanas (*Amblyrhynchus cristatus*). Essentially, a weaker negative feedback extends the stress response, and mimics a prolonged and sustained stressor such that the FCM concentrations are elevated into the reactive homeostasis range for a much longer period. As a result there the threshold between the reactive homeostasis range and homeostatic overload range progressively shrinks (Wright et al., 2023). For F1, this could imply that an additional stressor is likely to drive this animal into the homeostatic overload range and the pathological consequences associated with it. This obviously depends on duration and magnitude of the additional stressor (Romero et al., 2009). In comparison to the first two events, the stress response was elicited more immediately during the 3^rd^ visitor event across all joeys. It is possible that this could be due to the substantially shorter recuperation time between the 2^nd^ and 3^rd^ visitor event. Frequent repeated stressors do not allow enough time for the recovery of threshold between the reactive homeostasis range and homeostatic overload range (Wright et al., 2023). This implies that visitors that elicit an acute stress response from the koala joeys, if repeated frequently over several days (without recovery time) could drive the animal into the homeostatic overload range leading to chronic stress (unless the joeys get acclimatised to visitors). This again obviously depends on the rate of decline of the threshold between the homeostatic overload and the reactive homeostasis and whether or not joeys are acclimatised to visitors (Wright et al., 2023). We would like to reiterate that the discussion about the aforementioned scenarios are based on presumptions, and there is no direct evidence to confirm this. However, there are real-life examples [see (DuRant et al., 2016; Romero, 2012)] that incorporate the reactive scope model to justify (and predict) the physiological response to stressors with empirical evidence. Future research on koalas involving a variety of acute stressors may help get insights about the reactive scope model and the physiological coping mechanisms of koalas.

The concentration of FCMs in all koala joeys decreased within 1-3 days following visitor events, and remained within baseline until the next visitor event, indicating that the joeys were not chronically stressed. Instead, these results strongly suggest that the koala joeys are physiologically responding to environmental cues (visitors in this case) to maintain homeostasis. It is important not to overstate the impact of visitors observed in this study. It was evident that visitors were aversive enough to activate the HPA axis, but there was no direct evidence that this translated into a detrimental effect on the welfare of the joeys. In this study, all visitor encounters were acute, resulting in only transient elevations of FCM concentrations and there was no evidence of chronic stress in any animal.

## Conclusion

In summary, this study has provided a better understanding of the koala joey stress physiology and opened up new avenues for incorporating stress physiology models to comprehend the stress response of koalas. FCM profiles of koala joeys varied significantly and can be attributed to the natural variation between individuals. Daily profiling of FCM secretion revealed that visitors serve as an acute stressor for rehabilitating koala joeys and evoke a significant stress response. Whether or not koala joeys would elicit a similar response when encountered by familiar people (i.e. husbandry staff) remains to be investigated. The findings of this study have established a basis for the future monitoring of wellbeing in koala joeys and provide insight into how existing stress management strategies can be enhanced. It is important that visitor encounters be kept to a minimum and perhaps avoided altogether especially for young rehabilitating joeys. However, at the same time we also acknowledge that visitors, donors and the media, all contribute substantially to the overall broader goal of improving the conservation of koalas.

## Declarations

Not applicable

### Ethics approval and consent to participate

The study was conducted according to the guidelines of the Native or Exotic Wildlife and Marine Animals (NEWMA) Animal Ethics Committee and approved by the Institutional Review Board (or Ethics Committee) of The University of Queensland (approval number: AE000035).

### Consent for publication

Not applicable

### Availability of data and material

All data are contained within the article.

### Authors’ contributions

Conceptualization, E.N. and H.P.; methodology, E.N. and H.P.; formal analysis, E.N. and H.P.; investigation, E.N. and H.P.; resources, E.N. and H.P.; sample collection, A.B. and I.B. data curation, E.N. and H.P.; writing—original draft preparation, H.P.; writing—review and editing, E.N.; H.P.; A.B. and I.B.; supervision, E.N.; project administration, E.N. All authors have read and agreed to the published version of the manuscript.

## Acknowledgements

We thank the Magnetic Island Koala Hospital for their collaboration. We immensely appreciate the time, effort and the support offered by the staff in providing us with the samples and the veterinarians for providing identifying potential stressors encountered by the joeys.

## Abbreviations

FCM: Faecal cortisol metabolite;
EIA: Enzyme-immunoassay;
GC: Glucocorticoid;
HPA: Hypothalamo-pituitary adrenal;
HRP: Horseradish peroxidase;
ACTH: Adrenocorticotropic hormone;
RSM: Reactive Scope Model

